# Pleiotropic requirements for human TDP-43 in the regulation of cell and organelle homeostasis

**DOI:** 10.1101/560144

**Authors:** Agnes Roczniak-Ferguson, Shawn M. Ferguson

## Abstract

TDP-43 is an RNA-binding protein that forms cytoplasmic aggregates in multiple neurodegenerative diseases. Although the loss of normal TDP-43 functions likely contributes to disease pathogenesis, the cell biological consequences of human TDP-43 depletion are not well understood. We therefore generated human TDP-43 knockout cells and subjected them to parallel cell biological and transcriptomic analyses. These efforts yielded three important discoveries. First, complete loss of TDP-43 resulted in widespread morphological defects related to multiple organelles including: Golgi, endosomes, lysosomes, mitochondria and the nuclear envelope. Second, we identified a new role for TDP-43 in controlling mRNA splicing of Nup188 (nuclear pore protein). Third, analysis of multiple amyotrophic lateral sclerosis (ALS) causing TDP-43 mutations revealed a broad ability to support splicing of TDP-43 target genes. However, as some TDP-43 disease causing mutants failed to support the regulation of specific target transcripts, our results raise the possibility of mutation-specific loss-of-function contributions to disease pathology.

## Introduction

Transactivation response element DNA binding protein 43 (TARDBP, also known as TDP-43) belongs to the family of heterogeneous nuclear ribonucleoproteins (hnRNPs) (Purice and Taylor, 2018). Mutations in TDP-43 cause a familial form of amyotrophic lateral sclerosis (ALS) (Kabashi et al., 2008; Sreedharan et al., 2008) that is accompanied by the formation of neuronal cytoplasmic TDP-43 inclusions (Neumann et al., 2006). TDP-43 inclusions also occur in familial forms of ALS and frontotemporal dementia (FTD) that are caused by mutations in other genes as well as in sporadic forms of these and other neurodegenerative diseases (Amador-Ortiz et al., 2007; Ayaki et al., 2018; Ling et al., 2013; Mackenzie and Neumann, 2016; Rademakers et al., 2012). Cytoplasmic TDP-43 aggregates also occur in muscle in the context of inclusion body myopathy (Weihl et al., 2008).

The strong genetic and pathological links between TDP-43 and neurodegenerative disease have stimulated intense interest in elucidating the relationships between its normal and pathological functions (Taylor et al., 2016). Although TDP-43 was originally identified and named for its ability to bind to HIV-1 long terminal repeat DNA, it is now understood that TDP-43 is ubiquitously expressed in all cell types and plays an important physiological role in regulating the splicing of multiple endogenous human mRNAs (Appocher et al., 2017; Conlon and Manley, 2017; Ling et al., 2015; Tollervey et al., 2011). The specific RNA targets for TDP-43 vary between species. However, there is a conserved role for TDP-43 in suppressing the inclusion of cryptic exons via binding to UG dinucleotide repeats in their flanking regions (Chiang et al., 2010; Ling et al., 2015; Lukavsky et al., 2013; Polymenidou et al., 2011; Sephton et al., 2011; Tan et al., 2016). The loss of such activity results in the production of numerous frameshifted transcripts that are frequently targets of nonsense mediated decay. Identifying human genes affected by cryptic exon insertion arising from TDP-43 dysfunction and understanding the consequences of their disruption is thus important for understanding both the normal mechanisms whereby TDP-43 ensures splicing fidelity as well as the contributions of aberrant mRNA splicing to disease pathology. In addition to regulating mRNA splicing, TDP-43 has also been implicated in the regulation of other aspects of RNA biology including: transcription, microRNA processing, RNA stability and regulation of cytoplasmic ribonucleoprotein complexes such as stress granules, myo-granules involved in muscle regeneration and granules involved in axonal RNA transport in neurons (Gopal et al., 2017; Ratti and Buratti, 2016; Vogler et al., 2018).

Efforts to define TDP-43 function in mice through knockout (KO) strategies revealed that TDP-43 is absolutely required for embryonic development and viability (Chiang et al., 2010; Kraemer et al., 2010; Sephton et al., 2010; Wu et al., 2010). Even TDP-43 conditional KO strategies in specific cell types resulted in proliferation defects and/or cell death (Chiang et al., 2010). The lethality arising from TDP-43 depletion has limited efforts to define both normal TDP-43 functions as well as the cell biological consequences of TDP-43 depletion. As a result of these challenges, the disease contributions of nuclear TDP-43 depletion and/or TDP-43 inactivation associated with its cytoplasmic aggregation remain uncertain. Results from mouse studies are further complicated by the lack of conservation in TDP-43 targets between species (Ling et al., 2015; Prudencio et al., 2012). Studies in human cells where TDP-43 has been partially depleted (but not eliminated) by RNAi approaches have identified specific targets related to the functions of several organelles/pathways including autophagy and nuclear import (Ling et al., 2015; Prpar Mihevc et al., 2016; Stalekar et al., 2015; Xia et al., 2016). Although these results are intriguing, it remains unclear to what extent the regulation of any single TDP-43 target contributes to the total influence of TDP-43 on cell physiology.

As a comprehensive understanding of TDP-43 functions is critical for understanding normal human cell biology as well as for deciphering disease mechanisms, we have developed the first human TDP-43 knockout cells and used them to perform comprehensive cell biological and transcriptomic analyses of the consequences of TDP-43 depletion. The results of these experiments revealed that TDP-43 is required for the homeostasis of multiple subcellular organelles. Transcriptomic analysis of TDP-43 KO cells both confirmed the impact of TDP-43 on multiple known targets but also revealed new candidates. Given recent interest in the contributions of nuclear transport defects to neurodegenerative diseases associated with TDP-43 pathology (Chou et al., 2018; Gao et al., 2017; Kim and Taylor, 2017; Ward et al., 2014; Zhang et al., 2018), we highlight in particular the identification of the nucleoporin, Nup188, as a novel target of TDP-43-dependent splicing regulation. Furthermore, our analysis of the ability of multiple disease causing TDP-43 mutants to rescue TDP-43 KO phenotypes supports the general functionality of these proteins. These results suggest that loss-of-function arising from cytoplasmic accumulation and aggregation of mutant TDP-43 proteins rather than a direct loss of their fundamental splicing functions causes them to confer disease risk. However, we also identified TDP-43 mutations that are selectively defective in their ability to regulate specific target genes. This raises the interesting possibility that distinct properties of disease pathology might arise from these particular TDP-43 mutations.

## Results and Discussion

### Generation of a TDP-43 knockout cell culture model

Efforts to define TDP-43 function through genetic depletion approaches have previously been implemented in mice as well as in human cells in culture (Chiang et al., 2010; Kraemer et al., 2010; Ling et al., 2015; Prpar Mihevc et al., 2016; Sephton et al., 2010; Stalekar et al., 2015; Wu et al., 2010; Xia et al., 2016). Although these strategies have been informative in identifying many mRNA transcripts whose splicing depends on TDP-43 and physiological processes that are impaired in response to reduced TDP-43 levels, the essential functions of human TDP-43 have not been defined via a complete knockout. Furthermore, as the splicing targets of TDP-43 are highly species dependent, insights obtained from mouse studies have not always yielded results that were translatable to humans (Ling et al., 2015; Prudencio et al., 2012). These factors motivated us to generate human TDP-43 knockout cells via CRISPR-Cas9-mediated genome editing.

As past efforts to knockout TDP-43 reached the conclusion that this gene is essential and that its complete absence is not compatible with life, we first focused on HeLa cells to test the feasibility of a TDP-43 KO approach in human cells as this cell line is both highly adapted to robust growth in culture and very amenable to CRISPR-Cas9-mediated genome editing. Following transfection of HeLa cells with Cas9 and 2 distinct TDP-43-targeted small guide RNAs (sgRNAs) either alone or in combination, we analyzed polyclonal cell populations by immunoblotting and observed significant depletion of TDP-43 protein levels (Fig. 1A). Clonal populations of these cells were next isolated and screened by immunoblotting to identify clones that lacked the TDP-43 protein (Fig. 1B). The sgRNAs used for generating these KO cells target exon 3 (out of 8 exons in total) which encodes part of RNA Recognition Motif 1 (RRM1) near the beginning of the TDP-43 protein. Sequencing of this region of genomic DNA confirmed the loss of WT TDP-43 sequence at the Cas9 cut site in all copies of the TDP-43 gene and the presence of small insertions and deletions that result in frameshifts in our TDP-43 knockout cell line (Supplemental Fig. S1 A, B and C). In addition to prematurely terminating translation, the presence of frameshift mutations within this early exon are predicted to result in depletion of the TDP-43 transcript by nonsense mediated decay. Consistent with the conclusion that we successfully eliminated all production of TDP-43 protein, we did not detect any larger or smaller TDP-43 protein fragments that could have been generated by unexpected alternative splicing events that skip the mutated exon (Supplemental Fig. S1 D).

**Figure 1:**
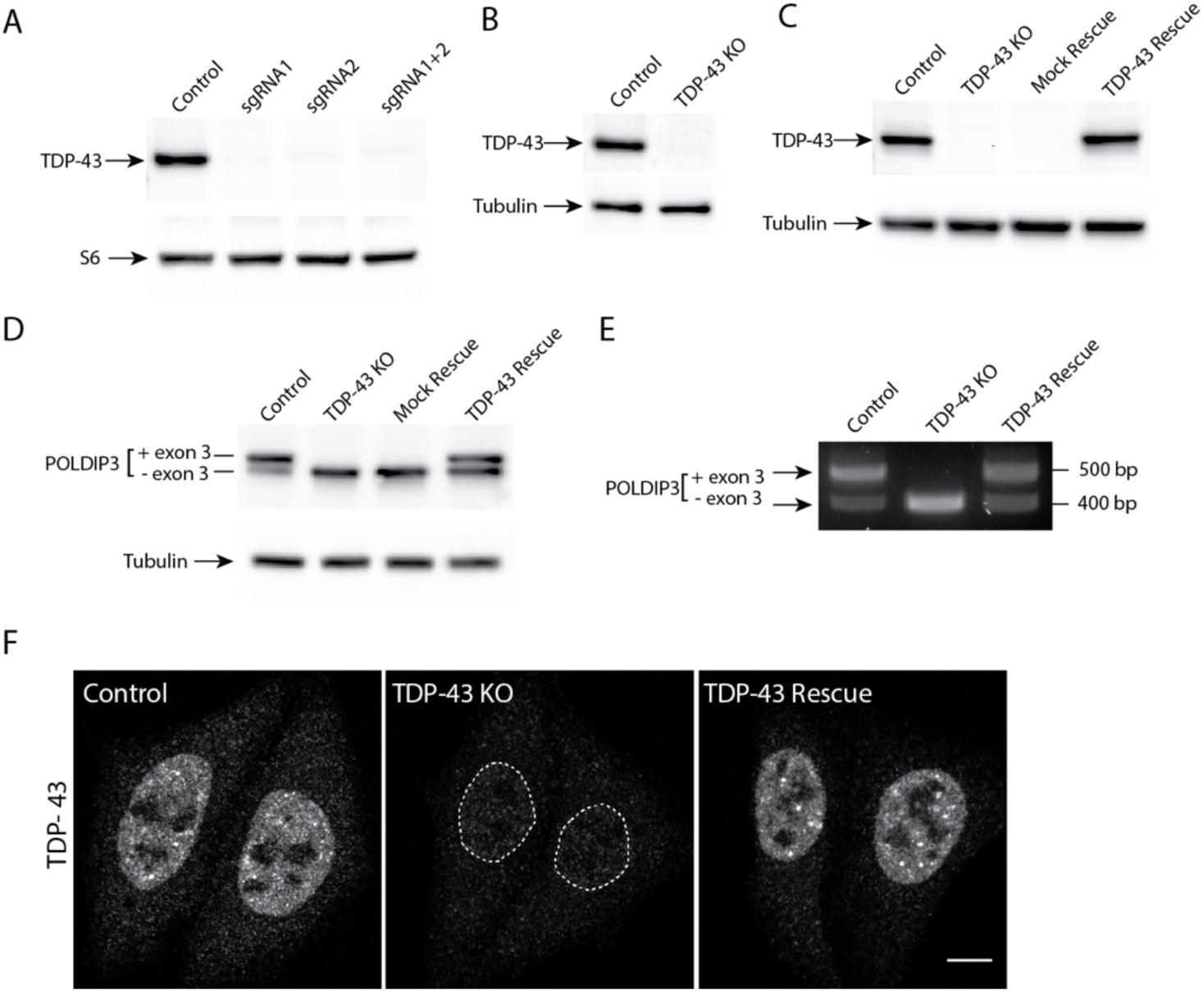
Generation and validation of human TDP-43 KO cells. **(A)** Immunoblot analysis of polyclonal populations of Hela cells transfected with Cas9 plus two sgRNAs against TDP-43 separately or in combination. Ribosomal protein S6 serves as a loading control. **(B)** Immunoblot analysis of a clonal population of TDP-43 KO HeLa cells. **(C)** Immunoblot evaluation of the restoration of TDP-43 expression via lentiviral transduction of TDP-43 KO cells with an untagged, wildtype form of TDP-43. **(D)** Immunoblotting revealed that levels of the larger POLDIP3 isoform were rescued by re-expression of TDP-43 in the TDP-43 KO cells. **(E)** Assessment of POLDIP3 exon 3 inclusion by RT-PCR. **(F)** Confocal immunofluorescence microscopy shows similar nuclear localization of the TDP-43 protein in both parental HeLa cells and the rescue cell line. Scale bar = 10µm.

The DNA sequencing and immunoblotting results both indicated that we had successfully generated human TDP-43 KO cells. To further characterize these cells, we examined POLDIP3, a well-defined target of TDP-43-dependent splicing regulation (Fiesel et al., 2012; Shiga et al., 2012). The POLDIP3 protein migrates as 2 bands on SDS-PAGE gels that reflect the presence or absence of exon 3, whose inclusion is dependent on TDP-43 (Fiesel et al., 2012; Shiga et al., 2012). In WT cells, the POLDIP3 protein migrated primarily at the expected size of 46 kDa with only a small amount of the smaller exon 3 lacking variant of this protein (Fig. 1D). In TDP-43 KO cells however, POLDIP3 migrated predominantly as the smaller form of the protein (Fig. 1D). The specificity of this TDP-43 KO phenotype was confirmed by stably reintroducing TDP-43 back into the KO cells via lentiviral transduction as this resulted in the restoration of WT levels of the TDP-43 protein (Fig. 1C) and a reversal of the POLDIP3 phenotype (Fig. 1D). TDP-43-dependent regulation of POLDIP3 mRNA splicing was further confirmed by RT-PCR analysis of RNA isolated from either WT, TDP43 KO or TDP-43 rescued KO cells (Fig. 1E). Immunofluorescence analysis identified a nuclear TDP-43 signal that was selectively present in only the WT and rescued cell lines while a minimal, non-specific, nuclear and cytoplasmic immunofluorescence signal was observed in the KO cells (Fig. 1F). These observations validate this antibody for future studies of TDP-43 subcellular localization.

It is striking that our rescue strategy employing lentiviral delivery of an untagged, human TDP-43 transgene driven by the strong cytomegalovirus (CMV) promoter yielded TDP-43 protein levels that so closely matched the endogenous expression levels of TDP-43. It is not clear whether this is simply fortuitous or whether it reflects a homeostatic mechanism whereby cells tightly control TDP-43 protein levels. Although TDP-43 auto-regulatory mechanisms have previously been described (Avendano-Vazquez et al., 2012; Ayala et al., 2011), our transgene lacks the key intronic and untranslated region (UTR) sequences that were previously shown to underlie known auto-regulatory mechanisms. Therefore, any homeostatic control of TDP-43 protein levels that might be occurring in our cells would have to arise via alternative mechanisms.

### Broad cell biological consequences of the TDP-43 KO

Previous studies of TDP-43 function in human cells have generally focused on widespread transcriptional changes or very specific phenotypes relating to a particular transcript and/or the organelle in which it functions (Ling et al., 2015; Prpar Mihevc et al., 2016; Stalekar et al., 2015; Xia et al., 2016). Although these approaches have yielded valuable insights, they left a gap with respect to our understanding of the full cellular impact of TDP-43 perturbations. We thus performed confocal microscopy analysis of cells that were stained in parallel with antibodies against established markers of multiple intracellular organelles. These efforts revealed a pleiotropic set of specific changes arising from the loss of TDP-43 that were reversed in the rescued cell line (Fig. 2). Key changes that were observed include: abnormal morphology of the nuclear envelope (Fig. 2A), Golgi fragmentation (Fig. 2B) early endosome dispersal (Fig. 2C) and clustering of both lysosomes (Fig. 2D) and mitochondria (Fig. 2E) in the perinuclear region.

**Figure 2:**
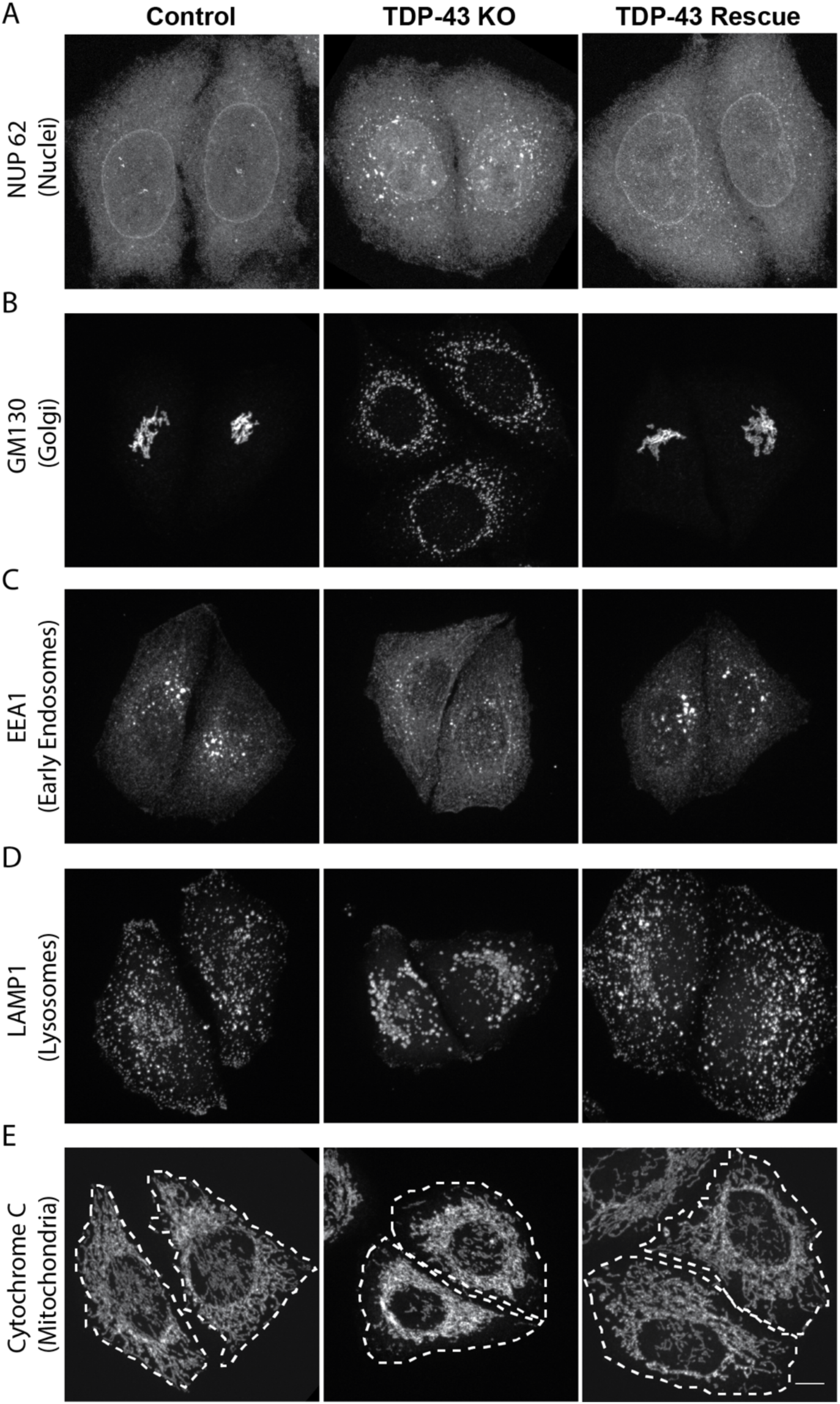
TDP-43 KO affects the morphology of multiple organelles. Representative confocal immunofluorescence images from cells of the indicated genotypes that were labeled with antibodies for the following organelles: **(A)** nuclear pore (Nup 62); **(B)** Golgi (GM130): **(C)** Early endosome (EEA1); **(D)** Late endosome/lysosome (LAMP1); **(E)** Mitochondria (cytochrome c). Scale bar = 10µm.

In addition to their abnormal subcellular distribution, the lysosomes in TDP-43 KO cells lacked cathepsin L (a major lysosomal protease that supports protein degradation, Fig. 3A). Cathepsin L localization to lysosomes was restored in TDP-43 rescue cells (Fig. 3A). Cathepsin L normally undergoes proteolytic processing upon delivery to lysosomes that is essential for its activation. This maturation of cathepsin L was impaired in TDP-43 KO cells and was restored in the TDP-43 rescue cells (Fig. 3D and E). Similarly to cathepsin L, cathepsin B processing in lysosomes of TDP-43 KO cells was also impaired (Fig. 3D). Cathepsin D on the other hand, still localized to the lysosomes in TDP-43 KO cells (Fig. 3B) suggesting that distinct mechanisms exist for the lysosomal delivery of these cathepsins. The maturation of the cathepsin D protein was likewise unaffected in TDP-43 KO cells (Fig. 3D).

**Figure 3:**
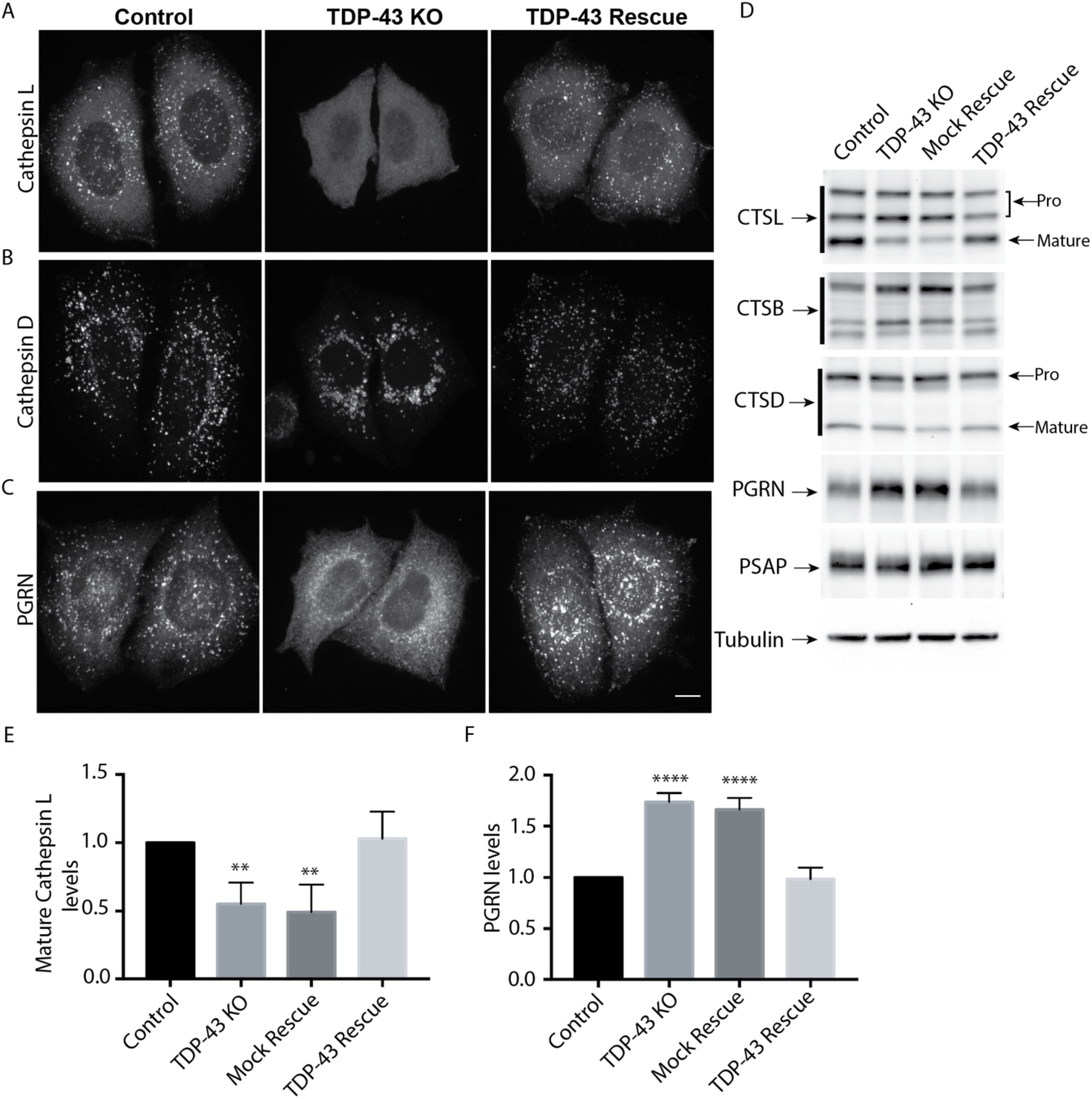
Characterization of lysosome-related defects in TDP-43 KO cells. **(A)** Representative confocal images of Cathepsin L localization in WT, TDP-43 KO and Rescued HeLa cell lines respectively. **(B)** Cathepsin D localization in cells of the indicated genotypes. **(C)** Impact of TDP-43 on PGRN subcellular localization. **(D)** Immunoblot analysis of lysosomal protein abundance and processing in cells of the indicated n=4, **p<0.0001, ANOVA with Bartlett post-test). Scale bar = 10µm.genotypes. **(E)** Quantification of the TDP-43-dependent changes in the abundance of the mature form of cathepsin L (n=4, p<0.005, ANOVA with Bartlett post-test). **(F)** Quantification of TDP-43-dependent changes in PGRN protein levels (n=4, **p<0.0001, ANOVA with Bartlett post-test). Scale bar = 10µm. Error bars show mean+/-SEM.

The lysosome defects in TDP-43 depleted cells were intriguing as mutations in the progranulin gene (GRN) give rise to FTD with TDP-43 pathology (Baker et al., 2006; Cruts et al., 2006) and the progranulin protein localizes to lysosomes (Gowrishankar et al., 2015; Hu et al., 2010). Interestingly, we observed that progranulin delivery to lysosomes was greatly diminished in the absence of TDP-43 (Fig. 3C). This was accompanied by a build-up of full length progranulin inside the cells (Fig. 3D and F), an effect that was restored to normal levels in TDP-43 rescue cells (Fig. 3D and F). Based on recent reports of efficient progranulin processing into granulin peptides within lysosomes (Holler et al., 2017), the increased abundance of full length progranulin is consistent with a defect in its lysosome delivery. Impaired function of cathepsin L, a major protease that mediates conversion of progranulin into granulins (Holler et al., 2017; Lee et al., 2017), could also contribute to this phenotype. In parallel with defects in lysosome localization and processing, we also observed greater secretion of the unprocessed forms of lysosomal proteins including: progranulin and cathepsins B, D and L (Supplemental Fig. 2). Interestingly, levels of prosaposin (PSAP), a protein that interacts with PGRN and directs its trafficking to lysosomes (Nicholson et al., 2016; Zhou et al., 2015), also exhibited more secretion in the TDP-43 KO cells (Supplemental Fig. 2).

Lysosome impairment arising from loss of TDP-43 is of potential interest in the context of neurodegenerative diseases such as ALS-FTD where human genetics and pathology have established roles for both TDP-43 and lysosomes as contributors to the disease process. This is particularly relevant in the case of GRN-linked FTD where GRN haploinsufficiency results in neurodegeneration that is accompanied by TDP-43 cytoplasmic aggregation (Gotzl et al., 2016; Mackenzie, 2007; Rademakers et al., 2012). Our observations of lysosome defects arising from TDP-43 depletion further suggest the possibility of a toxic feed forward loop wherein initially small losses of either TDP-43 or lysosome defects such as those arising from reduced progranulin levels could be amplified.

### TDP-43 is not essential for the formation of arsenite-induced stress granules

In addition to its splicing functions within the nucleus, TDP-43 is also found in cytoplasmic stress granules and has been implicated in their formation (Aulas et al., 2012; Freibaum et al., 2010; Liu-Yesucevitz et al., 2010). We observed that arsenite-induced stress granules containing T-cell-restricted intracellular antigen-1 (TIA1, an RNA binding protein) still formed in the absence of TDP-43 (Supplemental Fig. 3). Although we cannot rule out subtle defects in the kinetics of stress granule formation or effects on the formation of stress granules in response to other stimuli, this result indicates that TDP-43 is not absolutely required for stress granule formation.

### Transcriptomic analysis of TDP-43 KO cells

TDP-43 is well characterized for its ability to regulate mRNA splicing (Barmada, 2015; Gao et al., 2017; Ling et al., 2015). To gain a global view of mRNA changes arising from the absence of TDP-43, we performed RNA-Seq experiments wherein we compared the transcriptome of TDP-43 KO cells with those rescued via expression of WT TDP-43. The use of the rescued cells as our reference was supported by restoration of normal TDP-43 proteins levels as well as the rescue of normal POLDIP3 splicing in such cells (Fig. 1). Furthermore, since they were derived from the same KO clonal population, they control for any other adaptations or mutations that might have arisen in the course of generating the TDP-43 KO. Transcriptomic analysis of these 2 cell populations revealed numerous genes whose expression levels were significantly altered in response to the presence or absence of TDP-43 (Fig. 4A and Table S1). Featured prominently amongst the large group of proteins whose levels are decreased in the TDP-43 KO cells are established TDP-43 targets (Ling et al., 2015) such as ATG4B, INSR, PFKP and RANBP1. These changes in gene expression could arise due to direct effects of TDP-43 on the splicing of the affected mRNAs. For example, due to the inclusion/exclusion of exons resulting in frameshift mutations and degradation by nonsense mediated decay. However, indirect compensatory responses to the KO likely also contribute to the numerous more subtle changes that occurred in cells lacking TDP-43. We therefore next sought to more specifically identify genes whose splicing was altered in the TDP-43 KO cells (Fig. 4B, Table S2).

**Figure 4:**
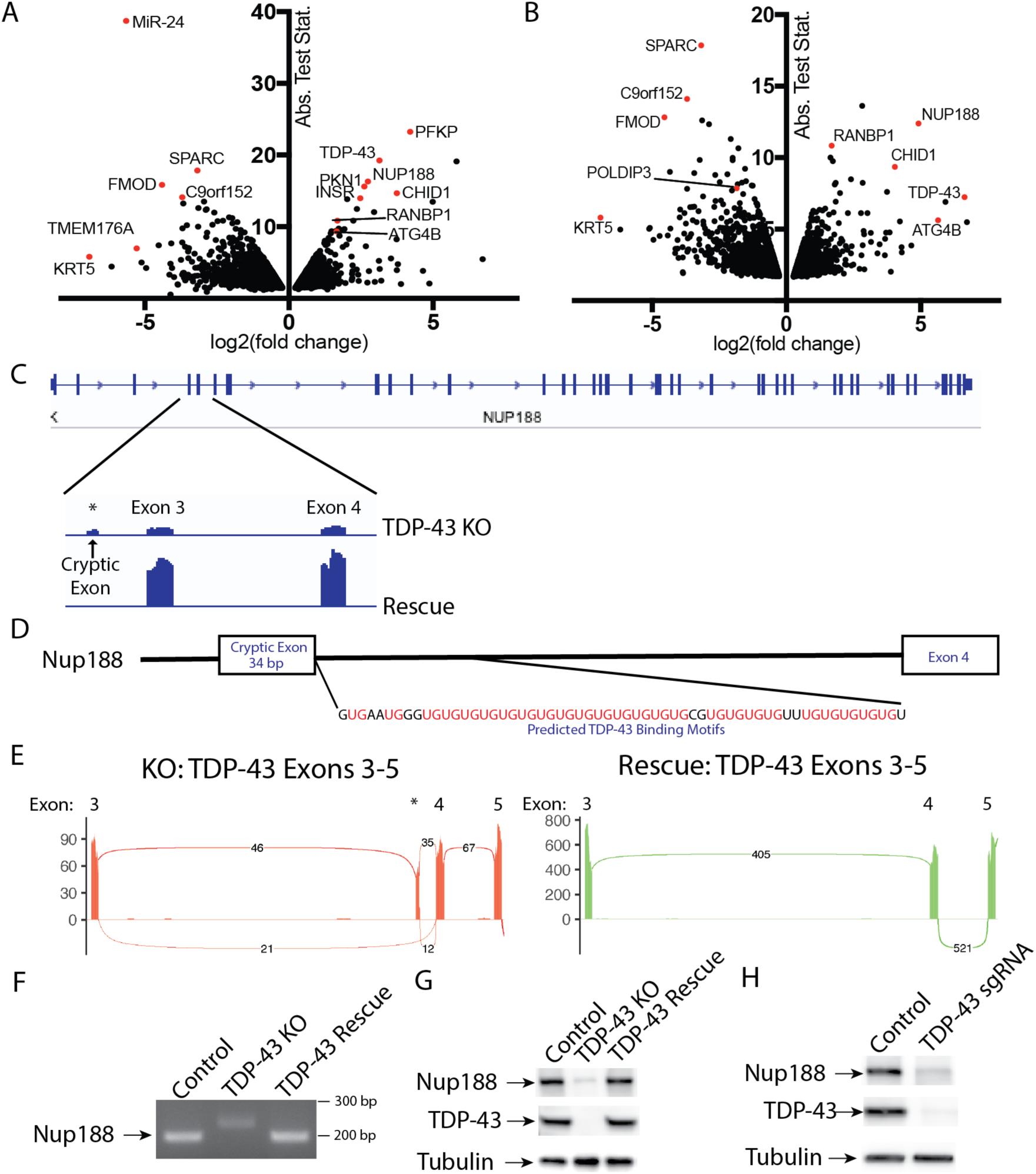
Identification of a cryptic exon in NUP188 as a TDP-43 target. **(A)** Changes in transcript abundance revealed by analysis of RNA-Seq data from TDP-43 KO versus Rescued cell lines. Fold change calculated for Rescue/KO HeLa cells. **(B)** Splicing changes detected by analysis of RNA-Seq data from TDP-43 KO versus Rescued cell lines. Fold change calculated for Rescue/KO cells. **(C)** Alignment of RNA-Seq reads showing the reduced overall abundance of Nup188 transcripts in TDP-43 KO cells as well as the increased abundance of a cryptic exon. **(D)** Presence of UG tandem repeats in the intron following the Nup188 cryptic exon. **(E)** Sashimi plots summarizing splicing patterns in the exon3 to exon 5 region of TDP-43 in KO and Rescue cell lines respectively. **(F)** RT-PCR validation of altered Nup188 splicing in TDP-43 KO cells. **(G)** Immunoblot analysis reveals the impact of TDP-43 on Nup188 protein levels. **(H)** Immunoblot analysis of Nup188 levels in Control versus TDP-43 depleted HEK293FT cells.

### Identification of Nup188 as novel TDP-43 target

Amongst the list of genes whose expression changed dramatically in a TDP-43-dependent manner was nucleoporin 188 (Nup188) whose levels fell by ∼6 fold in the absence of TDP-43 (Fig. 4F, Table S1). Analysis of differences in isoform abundance between the KO and Rescued cell lines revealed that this change in Nup188 expression in the TDP-43 KO cells was accompanied by the insertion of a cryptic 34 bp exon (Fig. 4C). In contrast, transcripts containing this cryptic exon are essentially undetectable in the TDP-43 expressing cells (Fig. 4C).

Closer analysis of the intronic sequence surrounding the cryptic exon in Nup188 predicts a direct role for TDP-43 in suppressing the inclusion of this exon. The TDP-43 RNA Recognition Motifs (RRMs) bind preferentially to RNA sequences containing tandem UG repeats and a hallmark of TDP-43 regulated splicing events is the presence of a UG-rich tract located 3’ of cryptic exons whose splicing is suppressed by TDP-43 (Ling et al., 2015; Lukavsky et al., 2013; Tollervey et al., 2011; Xiao et al., 2011). Analysis of this region in Nup188 reveals 25 UG pairs in the 60 base pairs immediately downstream of the 3’ splice site (Fig. 4D). The presence of this robust consensus TDP-43 binding site strongly suggests that Nup188 splicing is directly controlled by a canonical TDP-43-mediated mechanism. Further analysis of sequencing reads that span exon boundaries within this region clearly demonstrated the preferential inclusion of the cryptic exon in the TDP-43 KO cells and its complete absence when TDP-43 is present (Fig. 4E). This was further confirmed by RT-PCR analysis of amplicons spanning from exon 3 to exon 4 (Fig. 4F). Immunoblot experiments also detected a major reduction in Nup188 protein levels in the TDP-43 KO cells (Fig. 4G) that was rescued by re-expressing TDP-43. This major dependence of Nup188 splicing on TDP43 was independently replicated following CRISPR-mediated depletion of TDP-43 in HEK293FT cells (Fig. 4H).

As Nup188 plays an important role as a scaffold that contributes to nuclear pore assembly (Andersen et al., 2013), the reduction in Nup188 protein levels in TDP-43 KO cells likely contributes to the abnormal nuclear pore staining that we observed in these cells (Fig. 2A). However, the dysregulation of other previously reported TDP-43 targets implicated in nuclear pore function such as RANBP1 is likely to also contribute to this phenotype (Ling et al., 2015; Stalekar et al., 2015). Indeed, RANBP1 levels were also decreased in the TDP-43 KO cells in our transcriptomic analysis, although to a more modest degree than what was observed for Nup188 (Fig. 4A and Table S1).

### Cation independent mannose-6-Phosphate receptor (CI-M6PR) levels are reduced in TDP-43 KO cells

The impaired lysosomal delivery of multiple luminal proteins in TDP-43 KO cells raised questions about underlying mechanisms. Although many factors likely contribute to these lysosome-related phenotypes (including alterations that we observed in Golgi, endosome and lysosome morphology (Fig. 2B), we noted a ∼2x reduction in the abundance of CI-M6PR (also known as IGF2R) expression in TDP-43 KO cells (Table S1). CI-M6PR is a well characterized sorting receptor for the trans-Golgi network to endosome trafficking of multiple lysosomal hydrolases (Ghosh et al., 2003). Two factors motivated us to explore this change in more detail. First, to test whether even modest expression changes identified by RNA-Seq were indicative of detectable changes at the protein level. Second, to gain further insight into cellular changes that could contribute to the lysosome defects in TDP-43 KO cells. Consistent with the RNA-Seq results, we observed a parallel ∼2x decrease in the overall abundance of the CI-M6PR in the TDP-43 KO cells (Supplemental Fig. 4A and B). In contrast to the CI-M6PR, sortilin (SORT1), another sorting receptor involved in the trafficking of lysosomal hydrolases, did not show changes in transcript abundance, splicing or protein levels (Tables 1 and 2; Supplemental Fig. 4A) even though its splicing was previously reported to be regulated by TDP-43 (Prudencio et al., 2012). This may reflect known species specific preferences for the mouse SORT1 versus human SORT1 transcripts in the requirement for TDP-43 in regulating their splicing and/or cell type specific differences in this function of TDP-43 (Prudencio et al., 2012).

### Disease causing TDP-43 mutants rescue KO phenotypes but also reveal defects specific to individual mutations

Despite intense interest in elucidating the mechanisms whereby TDP-43 mutations give rise to neurodegenerative disease, it still remains unclear to what extent these mutations exert pathogenic effects via gain-of-function versus loss-of-function mechanisms (Kabashi et al., 2010; Orru et al., 2016; Vanden Broeck et al., 2015). We thus took advantage of robust TDP-43 KO phenotypes to assess the functionality of a panel of neurodegenerative disease causing TDP-43 mutants (Fig. 5A). To this end, we generated TDP-43 KO cell lines that stably expressed 8 different disease causing TDP-43 mutations (Fig. 5A; A90V, P112H, D169G, K263E, A315T, Q331K, M337V, A382T)(Banks et al., 2008; Corrado et al., 2009; Gitcho et al., 2008; Kabashi et al., 2010; Kabashi et al., 2008; Kovacs et al., 2009; Moreno et al., 2015; Rutherford et al., 2008; Sreedharan et al., 2008; Vanden Broeck et al., 2015; Winton et al., 2008). After confirming that each mutant was expressed at similar levels to the WT TDP-43 (Fig. 5B), we next examined their ability to rescue the expression levels/splicing of POLDIP3, PFKP [a previously characterized TDP-43 target gene whose splicing is altered in the elderly human brain (Ling et al., 2015; Raj et al., 2018)], and NUP188. Interestingly, even most TDP-43 mutants rescued the POLDIP, PFKP and Nup188 phenotypes, there were also specific instances where the rescue did not occur (Fig. 5B, C and D). This could at least partially reflect the fact that the suppression of cryptic exon inclusion mechanism whereby TDP-43 regulates Nup188 and PFKP is distinct from promoting exon retention in POLDIP3 (Fiesel et al., 2012; Ling et al., 2015). Consistent with the previous identification of an important role for RRM1 in supporting the regulation of POLDIP3 (also known as SKAR) splicing by TDP-43 (Fiesel et al., 2012), we observed that the P112H mutant rescued Nup188 and PFKP phenotypes but was deficient in restoring normal splicing of POLDIP3 (Fig. 5B, C, D, and E). However, as the K263E mutant was completely effective in rescuing Nup188 but not PFKP, additional factors must determine the functionality of specific TDP-43 mutant proteins. More broadly, these results indicate that in addition to loss of nuclear functions for TDP-43 mutants as they aggregate in the cytoplasm, individual TDP-43 mutants are also uniquely compromised in their ability to support the splicing of specific transcripts. These results stand in contrast to a previous report that disease-causing mutations have dominant effects on splicing (Arnold et al., 2013). This difference in conclusions could reflect the examination of physiological levels of mutant TDP-43 proteins in a KO background in our study versus over-expression on top of WT TDP-43 protein by Arnold et al. Although the panel of mutations that we examined did not reveal striking patterns between the regions where mutations resided and phenotypic consequences, the identification of differences in the functionality of these mutant proteins opens the door to future studies that would investigate in more detail whether mutations specific defects in splicing ability have any specific impact on variables of disease pathology such as the propensity to cause ALS versus FTD, age of onset and disease severity. More generally, our results establish the utility of this human TDP-43 knockout cell line as a valuable tool for the investigation of these and other disease causing TDP-43 mutations in the future.

**Figure 5:**
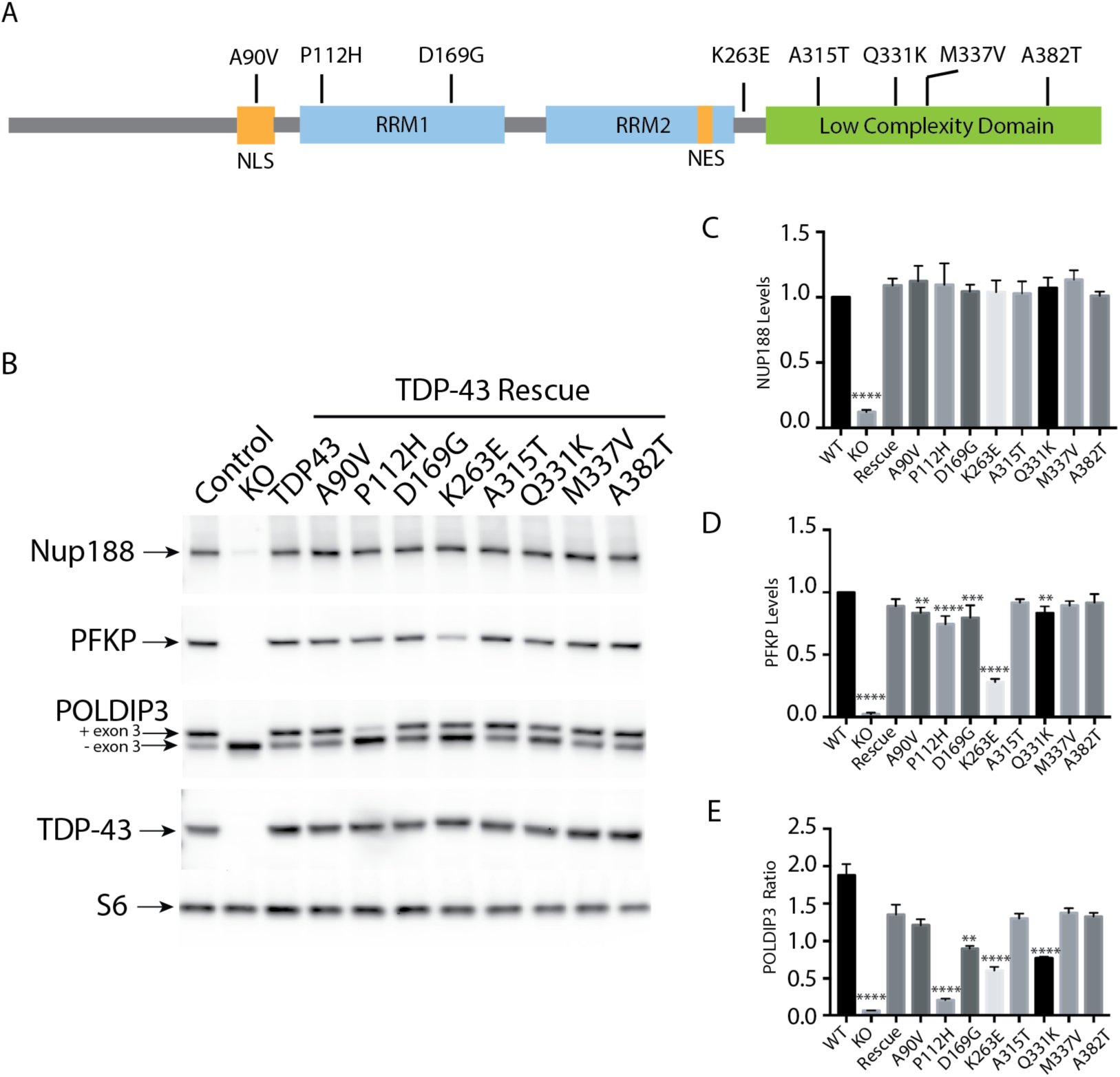
ALS-linked TDP-43 mutations selectively rescue the regulation of specific TDP-43 targets. **(A)** Schematic diagram that summarizes the TDP-43 protein domain organization and the location of mutants that were investigated in this study. **(B)** Representative immunoblots from Control, TDP-43 KO and TDP-43 KO HeLa cell lines that were reconstituted with TDP-43 variants containing the indicated ALS causing mutations. **(C)** Quantification of the ability of mutant forms of TDP-43 to rescue NUP188 abundance (n=3, ****p<0.0001, ANOVA with Dunnett’s multiple comparison test, comparison to WT). **(D)** Quantification of PFKP abundance (n=3, ****p<0.0001, ***p<0.0006, **p<0.0054, ANOVA with Dunnett’s multiple comparison test to compare mutant cell lines to WT). **(E)** Ratio of the POLDIP3 isoforms containing versus lacking exon 3 (n=3, ****p<0.0001, ***p<0.0003, **p<0.0016; ANOVA with Dunnett’s multiple comparison test to test for differences between the mutant cell lines versus WT rescue samples). Error bars show mean+/-SEM.

### Summary

Our focus on the essential functions of TDP-43 in HeLa cells was driven by the practical issues relating to viability that have precluded the complete KO of TDP-43 in other model systems. Although this strategy provides insights into the role for TDP-43 in the regulation of cellular processes that are common to all cell types, there are likely to be additional targets of TDP-43 that are unique to specialized cell types such as neurons, muscles and glia. Indeed, the pattern of splicing changes arising from TDP-43 depletion has been reported to vary between cell types (Jeong et al., 2017). Nonetheless, by taking advantage of a cell culture model that uniquely offered the opportunity to completely eliminate TDP-43 function, we have generated multiple new insights that broaden understanding of the pleiotropic cell biological consequences of eliminating TDP-43 expression and shed light on the functionality of disease causing TDP-43 mutants. Having observed that loss of TDP-43 affects multiple cellular organelles and hundreds of mRNA transcripts, it is challenging to attribute putative disease-causing mechanisms to the altered regulation of any one single transcript. However, due to the tremendous interest currently focused on defects in nuclear pore function in neurodegenerative disease associated with TDP-43 pathology (Chou et al., 2018; Kim and Taylor, 2017; Ward et al., 2014; Zhang et al., 2018; Zhang et al., 2015), it is interesting to consider how the NUP188 defects that we have uncovered in TDP-43 KO cells may intersect with other previously identified TDP-43-dependent components of the nucleo-cytoplasmic transport machinery such as RANBP1 (Ling et al., 2015; Stalekar et al., 2015). Our analysis of the ability of mutant forms of TDP-43 to rescue the splicing and expression levels of target genes in TDP-43 KO cells established that these cells are a useful tool for investigating the functionality of disease causing TDP-43 mutations. Our data furthermore raises new questions about whether subtle differences in TDP-43 protein functionality contributes to the pathology arising from distinct TDP-43 mutations.

## Materials and Methods

### Cell culture

HeLaM cells were kindly provided by Pietro De Camilli (Yale University) and were grown in Dulbecco’s modified Eagle’s medium (DMEM), 10% fetal bovine serum (FBS), and 1% penicillin/streptomycin supplement (all from Thermo Fisher Scientific). 293FT cells (Thermo Fisher Scientific) were grown on dishes coated with 0.1 mg/ml poly-d-lysine (Sigma) in the media described above. For stress granule induction, cells were incubated with 0.5 mM sodium arsenite (Fluka Analytical) prior to fixation.

### CRISPR/Cas9 genome editing

Use of the CRISPR/Cas9 genome-editing system to generate knock out cell lines was previously described (Amick et al., 2016). The guide RNA sequences used to generate TDP43 KO cell lines (summarized in Supplemental Table 3) were annealed and cloned into the BbsI site of pSpCas9(BB)-2A-Puro vector (px459, Feng Zhang, Addgene). Sub-confluent Hela cells were transfected with 1µg of px459 vector using FuGENE 6 (Promega). Transfected cells were selected with 2 µg/ml puromycin for 2 days and surviving cells were subsequently plated in 96 well plates at 1cell/well density to generate clonal lines. After selection and expansion of clonal populations, KOs were first identified by Western blotting and subsequently confirmed by sequencing of PCR-amplified genomic DNA (primers summarized in Supplemental Table 3). For HEK293FT experiments a polyclonal TDP-43 depleted population was examined following puromycin selection but without clonal selection.

### Lentiviral transduction

For rescue experiments, the human TDP-43 sequence was PCR-amplified from a previously described TDP43-tdTomato expression vector [kindly provided by Dr. Zuoshang Xu via Addgene (Yang et al., 2010), see table S3 for primer sequences]. The PCR product was introduced into the pLVX-puro vector (Clontech) by Gibson assembly (NEB). For ALS-associated TDP-43 mutations, cDNAs containing the mutations of interest were synthesized (gBlocks, IDT) and introduced into the pLVX-puro vector by Gibson Assembly (these plasmids will be made available through Addgene). For lentivirus production, 293FT cells (Invitrogen) were co-transfected with pLVX-TDP-43 along with the lentiviral packaging vectors pCMV-VSV-G (Addgene #8454), and pSPAX2 (Addgene #12260) in equal amounts using Fugene 6 transfection reagent. The lentivirus containing media was collected 24 hours post transfection, filtered through a 0.45 µm filter, combined with 8 µg/ml polybrene and added to the TDP-43 KO cell line. Transduced cells were selected with 2 µg/ml puromycin to establish polyclonal stable lines.

### Immunofluorescence and microscopy

Cells grown on 12-mm No. 1.5 coverslips (Carolina Biological Supply) were fixed with 4% paraformaldehyde (Electron Microscopy Sciences) washed with PBS and permeabilized with either 0.1% TX100 or 0.1% saponin in a 3%BSA/PBS buffer. Subsequent primary and secondary antibody incubations were carried out in the permeabilization buffer. Coverslips were mounted in ProLong Gold reagent supplemented with DAPI (Invitrogen). Images were acquired with either a Zeiss LSM 710 laser scanning confocal microscope with a 63x Plan Apo [numerical aperture (NA)=1.4] oil immersion objective and Zeiss Efficient Navigation software or an UltraVIEW VoX spinning disk confocal microscope (PerkinElmer) that consisted of a Nikon Ti-E Eclipse inverted microscope equipped with 60x CFI PlanApo VC, NA 1.4, oil immersion objective) and a CSU-X1 (Yokogawa) scan head that was driven by Volocity (PerkinElmer) software.

### Immunoblotting

Cells were lysed in Tris-buffered saline (TBS) supplemented with 1% Triton X-100 and protease/phosphatase inhibitor cocktails (Roche Diagnostics), the insoluble material was cleared by centrifugation for 10 min at 20,000 × *g*. Immunoblotting was performed by standard methods using 4-15% Mini-PROTEAN TGX precast polyacrylamide gels and nitrocellulose membranes (Bio-Rad). Ponceau S staining of membranes was routinely used to assess equal sample loading and transfer efficiency. Blocking was performed in 5% milk in TBS with 0.1% Tween 20. Primary antibody incubations were performed in 5% BSA in TBS with 0.1% Tween 20. Signals were detected with HRP-conjugated secondary antibodies (Cell Signaling Technology) and either Super Signal West Pico or Femto chemiluminescent detection reagents (Thermo Scientific) on a VersaDoc imaging system (Bio-Rad). ImageJ was used to measure band intensities. Antibodies used in this study are listed in Supplemental Table 4.

### RNA isolation and Reverse Transcription and PCR (RT-PCR)

Total RNA was extracted from WT, TDP-43 KO and rescue cell lines using the RNeasy kit (Qiagen). 400 ng of total RNA was reverse transcribed using the iScript cDNA Synthesis kit (Bio-Rad). Cryptic and alternatively spliced exons were PCR-amplified from the resulting cDNA with Q5 polymerase (NEB) and either Nup188 or POLDIP3 primers (IDT). The primers used to amplify the cryptic NUP188 exon were: 5’-ggagcagtagagaactgtgg-3’ and 5’-gctgattcttaaacccagttc-3’. The primers to amplify POLDIP3 splice variant were: 5’-gcttaatgccagaccgggagttg-3’ and 5’-tcatcttcatccaggtcatataaatt-3’. The resulting PCR reactions were resolved on 2.5% agarose gels supplemented with SYBR green (Invitrogen) and visualized on a VersaDoc imaging system (Bio-Rad).

### RNA-Seq

Six samples (3 pairs of biological replicates of TDP-43 KO and Rescue) were sequenced on Illumina’s HiSeq 2500 using 2×70bp paired-end reads, generating 104.4-128.2 million reads (52.2-64.1 million pairs) per sample. The sequencing reads were first trimmed for quality, trimming to the last base with quality >= 20 and dropping any read pair where either of the trimmed reads was shorter than 45 bp. The trimmed reads were aligned to the hg19 reference genome using TopHat2 (Trapnell et al., 2009), an RNA-seq aware aligner that allows for cross-exon, split alignments of reads to the genome. Overall, 96.8-97.1% of the paired-end reads aligned to the human genome, with an approx. 94.2-94.8% concordant pair alignment rate.

The Cufflinks and Cuffmerge software (Trapnell et al., 2013) was used to assemble the transcript isoforms from each sample’s reads and to merge the per-sample transcripts with the known human transcripts, to form a set of known and novel transcript isoforms from the data. This computation involves grouping overlapping reads together into “bundles”, constructing an “overlap graph” representing contiguously aligned regions and linked regions (where spliced alignments in a bundle link the aligned regions), then identifying the most parsimonious set of transcript isoforms that cover the overlap graph. Transcript abundance calculation and differential gene expression analysis, using the assembled known plus novel transcripts, was performed using Cuffdiff (Trapnell et al., 2013). Heatmaps and charts were visualized using Cummerbund (Trapnell et al., 2012), and Integrative Genomics Viewer (Thorvaldsdottir et al., 2013) was used for visualization of read depths and alignments. Sashimi plots were generated with ggsashimi using the alignment files generated with TopHat2 and hg19 annotation (Garrido-Martin et al., 2018).

## Acknowledgements

We are grateful to Patrick Lusk for reagents and advice related to analysis of the nuclear envelope. We appreciate support from the Yale Center for Genome Analysis for RNA-Seq experiments and from Sameet Mehta and James Knight for data analysis related to these experiments. Grants from the NIH (GM105718 and AG047270) and The Bluefield Project to SMF provided financial support for this research. The authors declare no competing financial interests.

## Supplemental Data

**Supplemental Figure 1:**
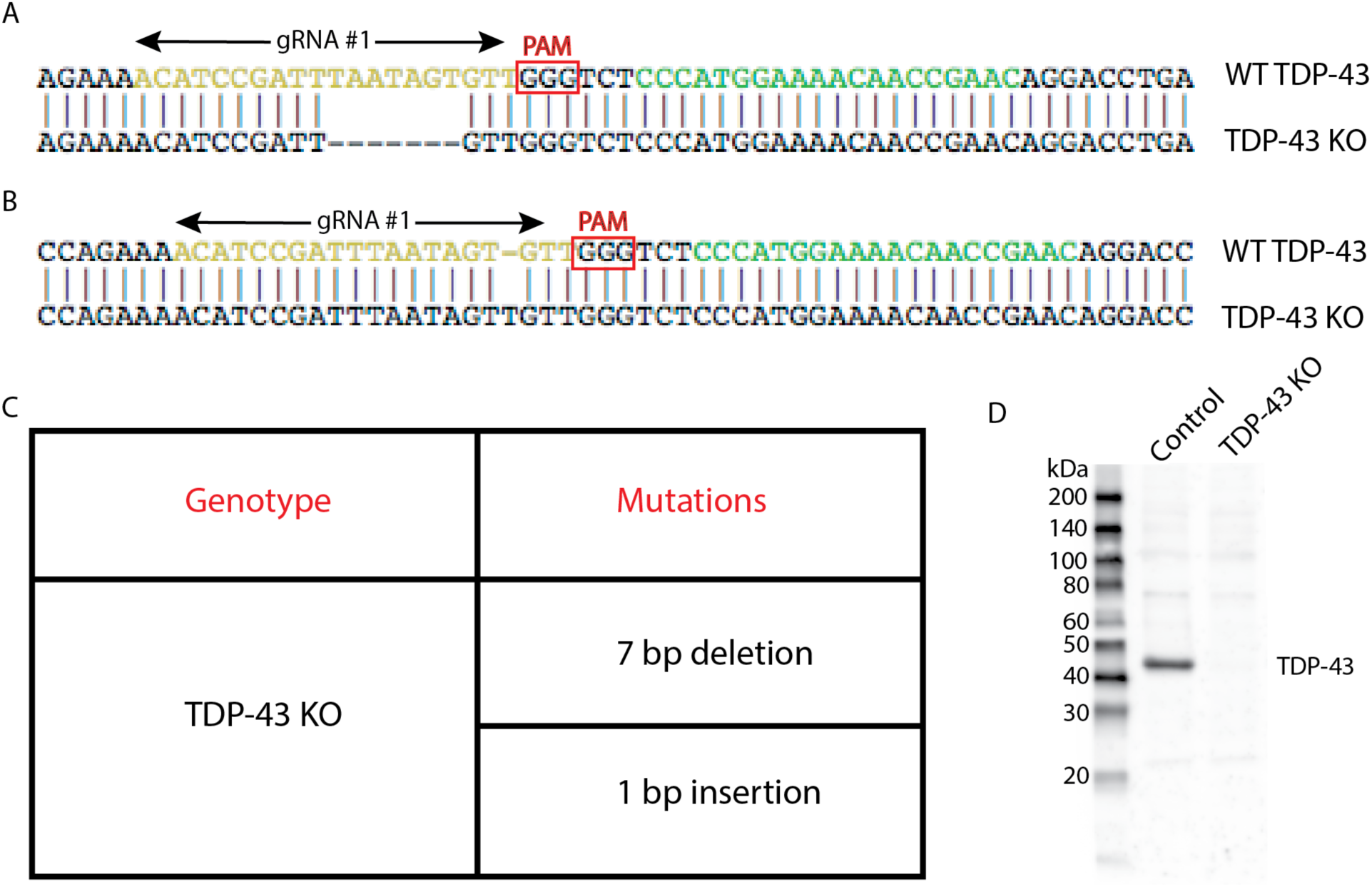
Generation of a TDP-43 KO cell line. **(A)** Alignment of WT sequence and mutant sequence containing a 7 bp deletion at the Cas9 cut site in TDP-43 exon 3 (region coding for RRM1). Regions recognized by gRNA#1 and gRNA#2 are highlighted in yellow and green respectively. Only gRNA#1 was used for the generation of theTDP-43 KO clonal line. **(B)** Alignment of WT sequence and mutant sequence containing a 1 bp insertion at the Cas9 cut site in TDP-43. **(C)** Summary of the mutations detected in the TDP43 KO cell line by sequencing of the region of genomic DNA targeted by TDP-43 sgRNA. **(D)** Immunoblotting results from WT and TDP-43 KO cells reveal the absence of both full length TDP-43 protein as well as of any fragments arising from the mutant alleles.

**Supplemental Figure 2:**
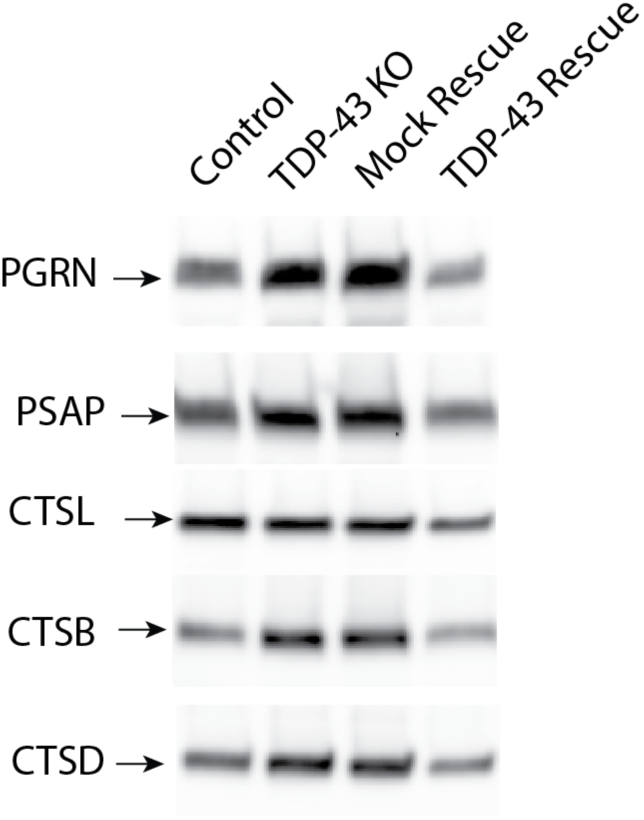
Immunoblot analysis of lysosome proteins secreted into the media by cells of the indicated genotypes. These immunoblots all show the “pro” form of the respective lysosomal proteins.

**Supplemental Figure 3:**
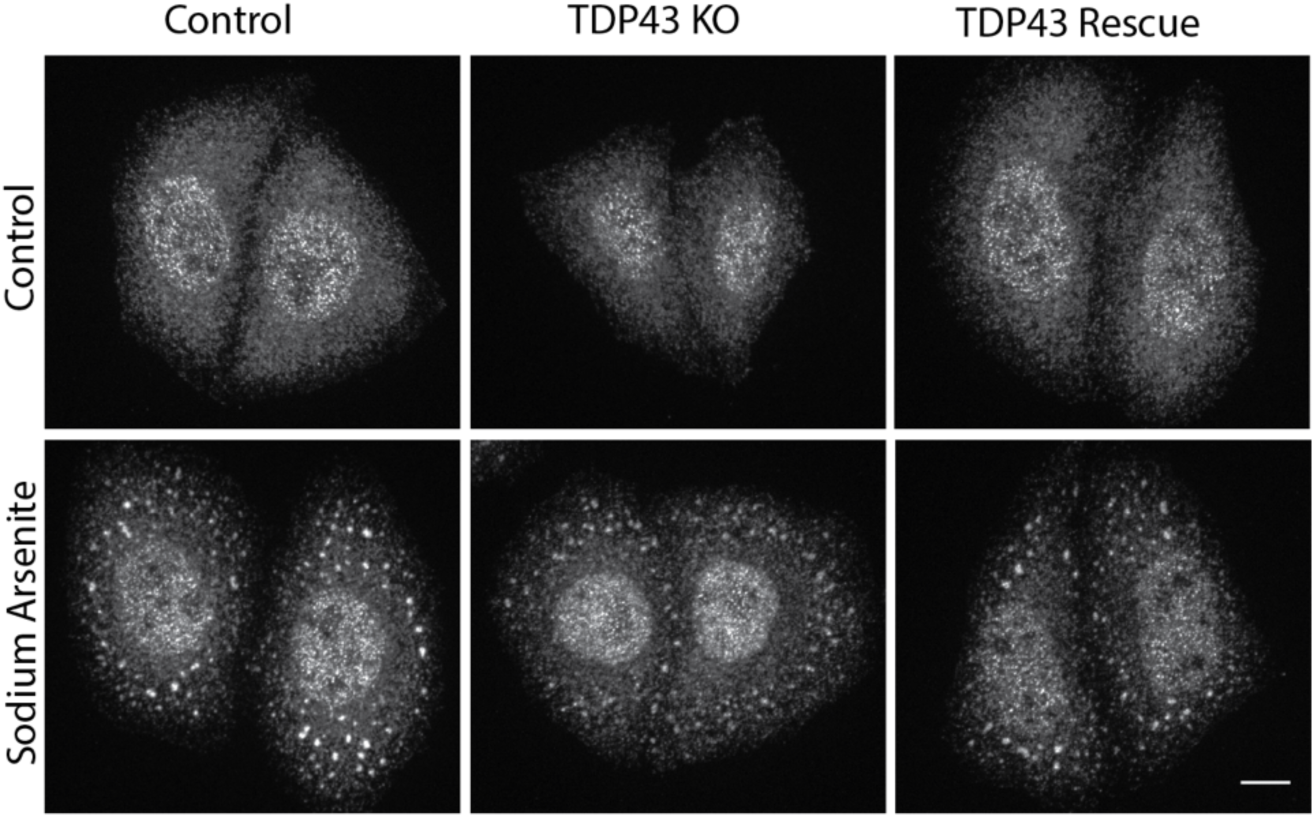
TDP-43 is not required for the formation of arsenite-induced stress granules. Immunofluorescent analysis revealed that TIA-positive stress granules form in TDP-43 KO cells in response to sodium arsenite treatment (0.5 mM, 15 min).

**Supplemental Figure 4:**
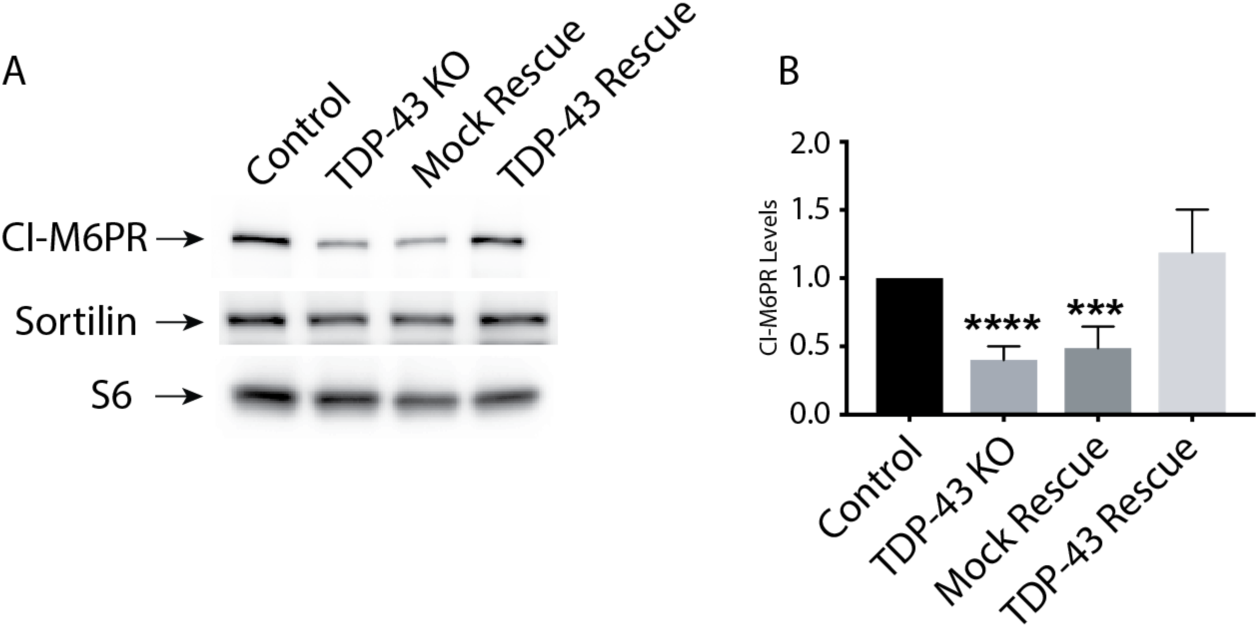
Impact of TDP-43 KO on CI-M6PR and SORT1 protein levels. **(A)** Representative immunoblots demonstrating that TDP-43 KO cells exhibit a reduction in CI-M6PR levels while SORT1 levels are unaffected. **(B)** Quantification of CI-M6PR protein levels in cells of the indicated genotypes (n=4, ****p<0.0001, ***p=0.0001, ANOVA with Bonferroni post-test). Mean+/-SEM.

**Supplemental Table 1:** Summary of gene expression changes detected in RNA-Seq analysis of TDP-43 KO and Rescue cell lines. Data is presented in the accompanying Excel file.

**Supplemental Table 2:** Summary of isoform expression/splicing changes detected in RNA-Seq analysis of TDP-43 KO and Rescue cell lines. Data is presented in the accompanying Excel file.

**Supplemental Table 3:**
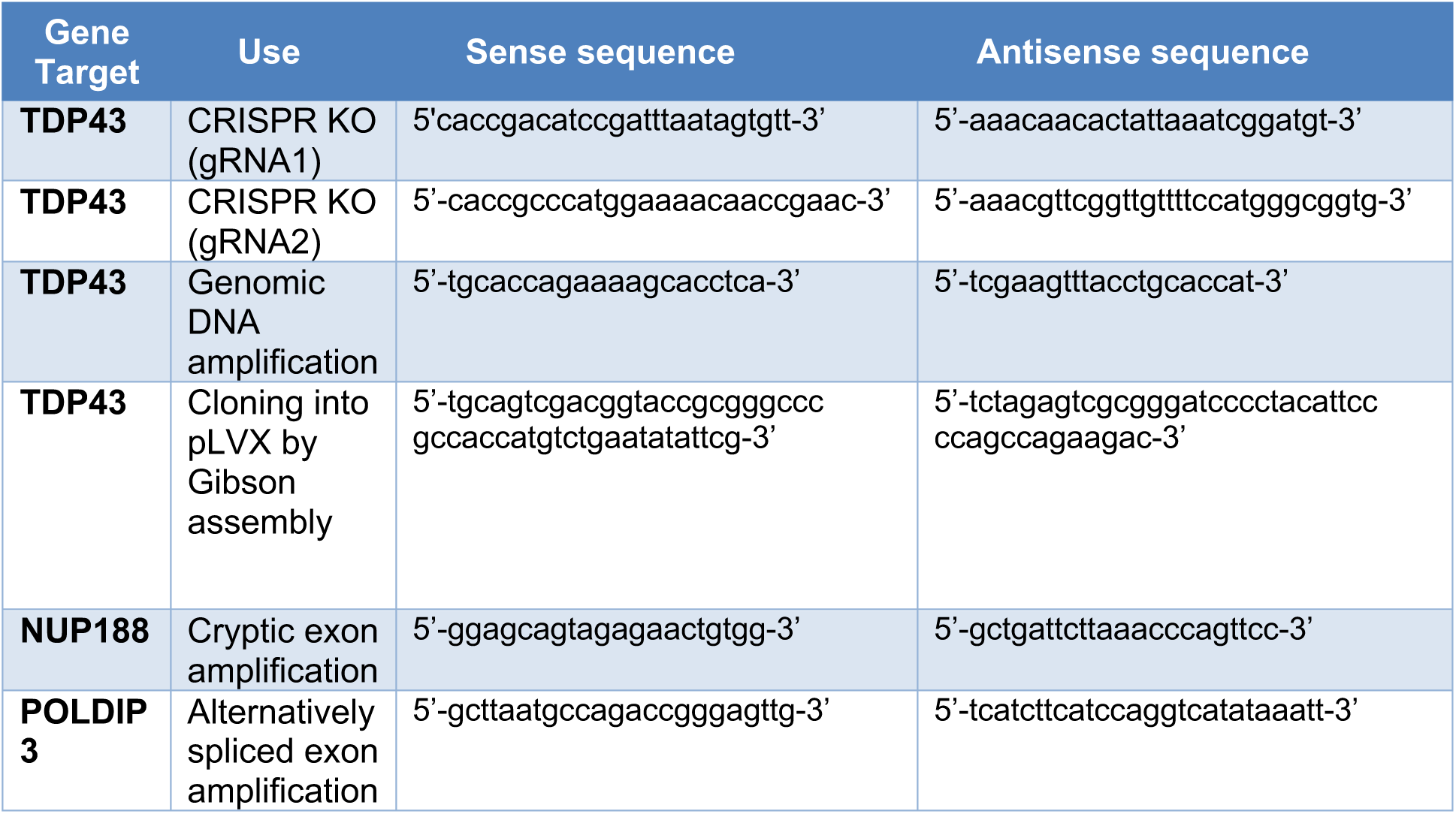
Summary of oligonucleotide primers used in this study.

**Supplemental Table 4:** Summary of antibodies used in this study.

| **Antibody** | **Source** | **Catalog/Clone number** |
| --- | --- | --- |
| **Cathepsin D** | R&D Systems | AF1014 |
| **Cathepsin L** | R&D Systems | AF 952 |
| **Cytochrome C** | BD Pharmingen | 556432 |
| **EEA1** | Thermo Fischer Scientific | PA1-063A |
| **GM130** | BD Transduction Laboratories | 610822 |
| **LAMP1** | University of Iowa DSHB | H4A3 |
| **Nucleoporin p62** | BD Biosciences | 610497 |
| **Nup188** | Bethyl Laboratories | A302-322A |
| **PFKP** | Cell Signaling | 5412S |
| **PGRN** | R&D Systems | AF2420 |
| **POLDIP3 (SKAR)** | Proteintech | 17466-1-AP |
| **Saposin C** | Santa Cruz Biotechnology | sc-32875 |
| **Ribosomal Protein S6** | Cell Signaling Technology | 2217 |
| **TDP43** | Cell Signaling Technology | 3449S |
| **TIAR** | Cell Signaling Technology | 5137S |
| **Tubulin** | Sigma -Aldrich | T5168 |

